# Correlative Light Electron Ion Microscopy reveal *in vivo* localisation of bedaquiline in *Mycobacterium tuberculosis* infected lungs

**DOI:** 10.1101/2020.08.05.237537

**Authors:** Antony Fearns, Daniel J. Greenwood, Angela Rodgers, Haibo Jiang, Maximiliano G. Gutierrez

## Abstract

Correlative light, electron and ion microscopy (CLEIM) offers huge potential to track the intracellular fate of antibiotics, with organelle-level resolution. However, a correlative approach that enables subcellular antibiotic visualisation in pathogen-infected tissue is lacking. Here, we developed CLEIM in tissue (CLEIMiT), and used it to identify the cell-type specific accumulation of an antibiotic in lung lesions of mice infected with *Mycobacterium tuberculosis.* Using CLEIMiT, we found that the anti-TB drug bedaquiline is localised not only in foamy macrophages in the lungs during infection but also accumulate in polymorphonuclear (PMN) cells.

## Introduction

An effective chemotherapy against bacterial infections must include antibiotics with pharmacokinetic properties that together allow penetration into all infected microenvironments[1]. Antimicrobial penetration is especially important for the treatment of infections where antibiotics need to reach intracellular bacteria[2], including *Mycobacterium tuberculosis*. In tuberculosis, treatment requires at least three antibiotics for six months[3], and we do not fully understand why this extended treatment is needed. In this context, understanding how tissue environments affect antibiotic localisation, exposure, and consequently efficacy against the pathogen, is crucial [4].

Although it is critical to define if antimicrobials are able to reach their intracellular targets, imaging of antibiotics (and drugs in general) at the subcellular level in infected tissues remains challenging. Only recently have studies *in vivo* determined antibiotic distributions in granulomatous lesions by matrix-assisted laser desorption-ionisation mass spectrometric imaging (MALDI-MSI)[5]. However, this approach only allows analysis at the tissue level and does not reach subcellular or even cellular resolution[6]. On the other hand, nanoscale secondary ion mass spectrometry (nanoSIMS) has been used to visualise drugs at 50 nm resolution in cells [7] and tissues [8]. However, there are limitations with this method such as the lack of correlation with other available imaging modalities that provide spatial information of specific cell types localisation and function. Thus, correlative approaches are needed to obtain both spatial localisation of drugs and biologically relevant information from experimental systems[9]. Recently, a correlative imaging approach combining light, electron and ion microscopy (CLEIM) has been developed for subcellular antibiotic visualisation *in vitro* cultured cells [10]. However, there are currently no approaches available that allow correlative studies at the subcellular resolution *in vivo*.

## Results

With the aim to define the subcellular localisation of antibiotics in infected cells within tissues, we used a mouse model of tuberculosis. Our goal was to develop an imaging approach to analyse the distribution of antibiotics from complex tissues to individual cells at the subcellular level in infected lungs. For that, we infected susceptible C3HeB/FeJ mice with *Mycobacterium tuberculosis* H37Rv expressing fluorescent E2-Crimson via aerosol infection (**Figure 1A**). The C3HeB/FeJ susceptible mouse strain develops necrotic lesions in the lung that better recapitulate human granulomas, a hallmark of tuberculosis infection[11]. After 21 days of infection, mice were treated daily for five days either with control vehicle or 25 mg/kg of the anti-mycobacterial antibiotic bedaquiline (BDQ). As previously reported[12], this treatment reduced approximately ten-fold the bacterial loads in the lungs, as measured by colony forming units (CFU) counting (**Figure S1A/B**). Following treatment, mice were euthanised and fixed by perfusion with formalin. Lungs were removed and granulomatous lesions were visualised by Micro Computed Tomography (μCT, **Figure 1A, Figure S1C and Movie S1**). Replicate lung tissues were embedded in agarose for further processing and imaging.

**Figure 1.**
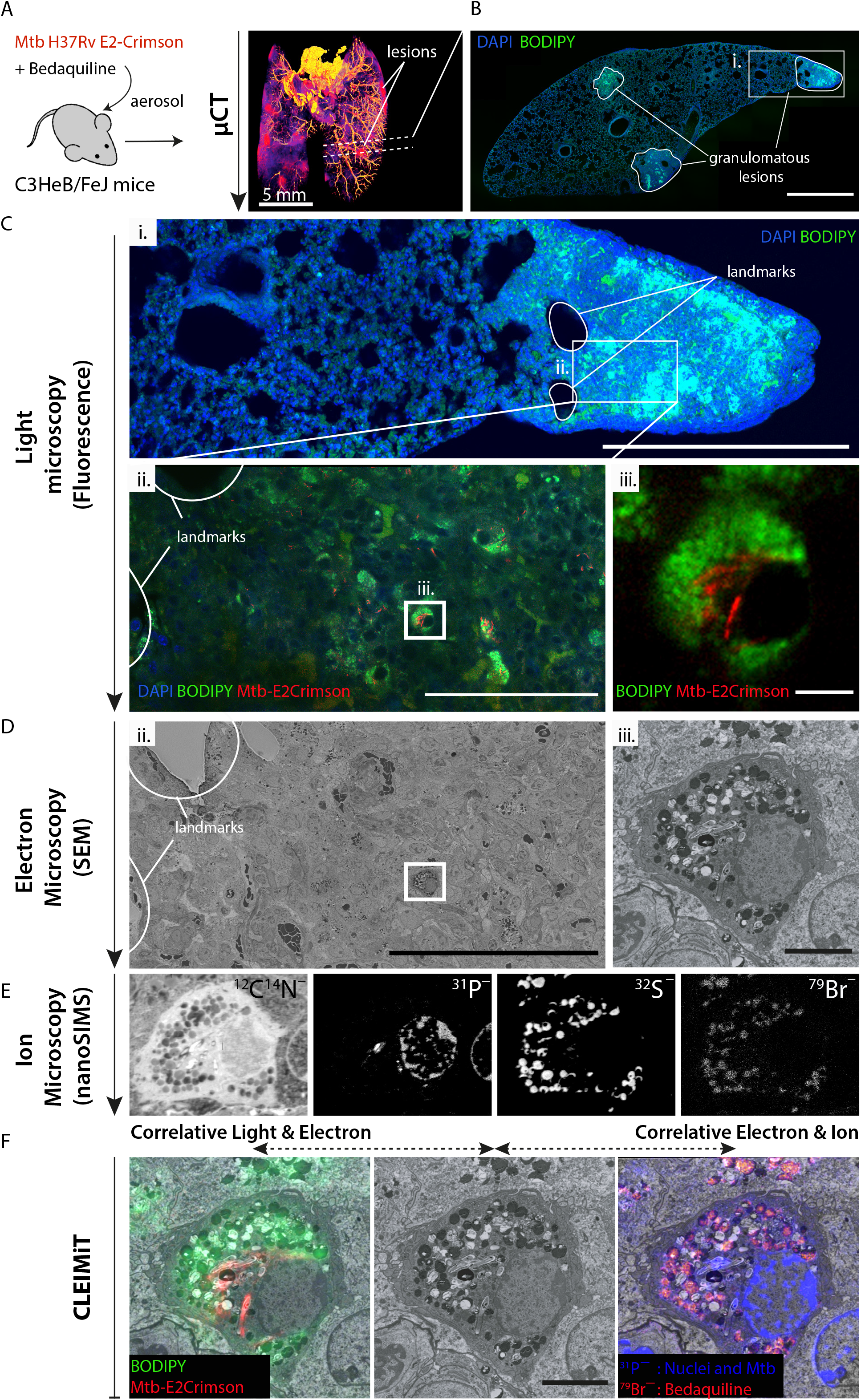
The CLEIMiT workflow and correlative imaging of BDQ in *M. tuberculosis*-infected foamy macrophages within lung tissue. **(A)** Diagram illustrating the *in vivo* experimental setting. C3HeB/FeJ mice were infected with *Mycobacterium tuberculosis* H37Rv expressing E2-Crimson (Mtb-E2Crimson) by aerosol. After 21 days, infected mice were treated with 25mg/kg of Bedaquiline (BDQ) or vehicle daily for 5 days via oral gavage. Lungs were removed, fixed with 10% formalin, contrasted and embedded in low melting point agarose then imaged by μCT for sequential vibratome sectioning. **(B)** Fluorescence microscopy: A tile scan of a tissue section (~100 μm thickness) stained with DAPI (blue) and BODIPY (green), granulomatous lesions are marked with a solid white line to indicate the boundary (scale bar = 1000 μm). **(C)** Light microscopy of a Region of Interest (ROI): (i) zoomed fluorescence image (white box from figure 1B), “landmarks” used for downstream location recognition, are indicated by the solid white boundary lines (scale bar = 500 μm). white rectangle shows the ROI for downstream analysis. (ii) A confocal image of the region indicated by the white box above shows an area of strong cellular infiltration and the accumulation of BODIPY (green) positive cells. Cells infected with Mtb-E2Crimson (red) are also visible throughout this region. The same landmarks marked in image (i) are present. white box indicates the selected infected foamy cell. Scale bar = 100 μm. (iii) Zoomed in image showing the selected foamy cell infected with Mtb-E2Crimson (red) for correlative analysis. Scale bar = 5 μm. **(D)** Electron microscopy of the ROI: (ii) tissue overview (600x magnification) with landmarks present, white box indicates the selected infected foamy cell. Scale bar = 100 μm. (iii) Zoomed in image showing the selected foamy cell infected with Mtb (15,000x magnification). Scale bar = 5 μm. **(E)** Ion microscopy of the selected cell: Panel shows the individual nanoSIMS images for the following ion signals from left to right; ^12^C^14^N^−^, ^31^P^−^, ^32^S^−^ and ^79^Br^−^. **(F)** Correlative light, electron and ion microscopy in tissue (CLEIMiT): Left, a correlated image overlaying fluorescent signal from BODIPY (green) and Mtb-E2Crimson (red) against the SEM image. Right, a correlated image overlaying the ^79^Br^−^ and ^31^P^−^ signals with the SEM image. Center, the corresponding SEM image of the infected foamy cell. Scale bar = 5 μm.

One of the main technical challenges of our attempt to define if the antibiotic reached intracellular bacteria was to identify and correlate across the different imaging modalities and scales the infected cells present in the lung. We devised a strategy that included the identification of a granulomatous lesion within 100 μm thickness sections and non-destructive 3D imaging by confocal laser scanning microscopy of the entire section as well as the region of interest (**Figure 1B and 1C**). Vibratome sections were stained with DAPI to visualise nuclei and BODIPY 493/503 to visualise lipid droplets (LD), previously shown to accumulate in foamy macrophages in necrotic lesions[13]. In agreement with previous studies, we found that granulomatous lesions were heavily enriched in LD-laden foamy macrophages (**Figure 1B and 1C**). After fluorescence imaging, sections were recovered and resin embedded for electron microscopy. In order to correlate the 3D fluorescence microscopy with the electron microscopy, sections were analysed by μCT 3D scanning (**Figure S2**). This approach enabled the precise localisation of the ROI previously imaged by fluorescence, and the angle correction during sectioning (**Figure S2**). In this way, the section obtained for Scanning Electron Microscopy (SEM) and nanoSIMS could be matched to the 3D fluorescence image with a high degree of accuracy (**Figure S2**). The sections were then imaged by SEM (**Figure 1D**) and the same section was then coated with 5 nm gold and transferred for nanoSIMS analysis. BDQ contains a bromine atom, so we determined its localisation by the intensity of the ^79^Br ion signal[10]. The regions imaged by SEM were identified using the optical microscope in the nanoSIMS. The sample was scanned with a focused ^133^Cs^+^ and secondary ions (^12^C^−^, ^12^C^14^N^−^, ^79^Br^−^, ^32^S^−^ and ^31^P^−^) and secondary electrons were collected (**Figure 1E**). The ^12^C^14^N^−^ and ^31^P^−^signals were useful to show the morphology of cells and tissues, with ^12^C^14^N^−^ signals largely from proteins and the highest ^31^P^−^ signals are from nucleic acids and structures we believe are polyphosphates in Mtb.

To correlate across imaging modalities with subcellular resolution, endogenous structures were used as landmarks. LD were located by fluorescent staining in the optical image, ultrastructure in the SEM image, and ^32^S^−^ signal in the ion image. The ^32^S^−^ signal was due to the osmium/thiocarbohydrazide staining of lipids. Bacteria were localized by fluorescence (E2-Crimson signal), ultrastructure and ^31^P^−^ signal in the ion image. The cell nucleus was aligned using ultrastructure and the ^31^P^−^ signal (**Figure 1F**). Concurrent with previous CLEIM *in vitro* studies, we found that BDQ accumulated heterogeneously in LD and Mtb, with particularly high levels in infected foamy macrophages (**Figure 1F and Figure S3**). Importantly, some bacteria contained high levels of the antibiotic whereas others did not show any signal, indicating that the antibiotic is not able to evenly reach throughout intracellular bacteria present in the infected tissue (**Figure 1F**).

Taking advantage of this method, we then focused on a more quantitative approach (**Figure S4**) to analyse intracellular antibiotic localisation in the lung lesions. For that, we performed a combined tile scanning by SEM and nanoSIMS covering larger areas of the tissue (**Figure 2A**). This allowed to define the distribution of BDQ in single cells and bacteria (**Movie S2**). Unexpectedly, we found that BDQ not only localised in lipophilic environments (*e.g.* in LD) but also in non-lipophilic cellular environments. Specifically, we found that BDQ strongly accumulated in polymorphonuclear cells (PMN). Antibiotic-rich PMN were present both alongside (**Figure 2B-C**) and away from areas enriched with foamy macrophages (**Figure 2D**). In contrast to macrophages where the ^79^Br^−^ signal was primarily associated with LD; in PMN, the ^79^Br^−^ signal was not only associated with granules but also with the cytosol (**Figure S5**). Thus, in tissues, BDQ accumulates in a cell-type dependent manner across two cell populations (foamy macrophages and PMN) with very different metabolic and functional properties. Confirming our previous observations, quantitative analysis revealed that BDQ heterogeneously accumulated in LD and Mtb (**Figure 2E**). Both Mtb outside and inside LD accumulated BDQ (**Figure 2E**). These PMN cells are likely neutrophils recruited to the granuloma as reported in this mouse model of TB infection[14]. Neutrophils are rich in Myeloperoxidase (MPO), a peroxidase that produces hypochlorous acid from hydrogen peroxide and chloride anion or hypobromous acid if bromide anion is present[15]. However, in untreated mice the ^79^Br^−^ signal was significantly lower and only slightly associated with PMN granules, indicating that the ^79^Br^−^ signal was primarily coming from the antibiotic (**Figure S5**). ^79^Br^−^/^12^C^14^N^−^ were used when comparing the BDQ-treated and non-treated tissues. The normalisation to ^12^C^14^N^−^ was to compensate possible minor variations in the primary ion current during imaging.

**Figure 2.**
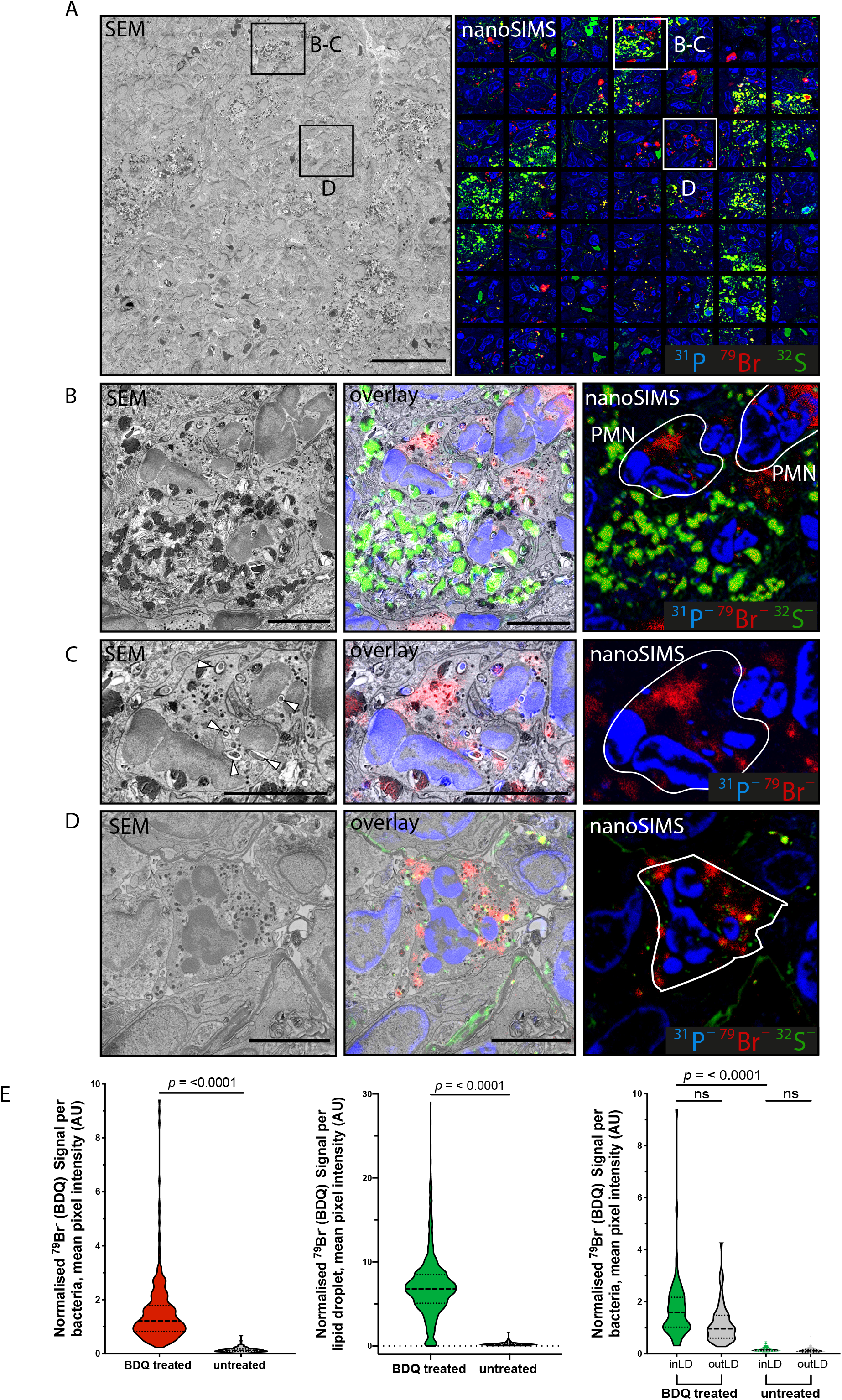
Quantitative distribution of BDQ reveals intracellular distribution of BDQ in foamy macrophages and PMN within granulomatous lesions. **(A)** Left, a tiled SEM image of a region of granulomatous lung tissue indicating the zoomed areas. Scale bar = 40 μm. Right, a mosaic of 49 individual ion micrographs showing the quantitative distribution of ion signals in the respective area of tissue. Sulphur ^32^S^−^ is shown in green, bromine ^79^Br^−^ in red and phosphorus ^31^P^−^ in blue. **(B)** Polymorphonuclear cells (PMN) are recruited to the foamy macrophage rich lesions. SEM of zoom 1 (left) and the distribution of secondary ions ^31^P^−^, ^79^Br^−^ and ^32^S^−^ (right). Overlaid image between SEM (left) and Secondary ion (right) is shown in the center. PMN are demarcated by the white boundary line. Scale bar = 5 μm **(C)** An infected PMN from panel B showing strong accumulation of BDQ. Overlay between the SEM and secondary ion signals for ^79^Br^−^ (red) and ^31^P^−^ (blue) is depicted in the center. White arrowheads indicate intracellular bacteria. Scale bar = 5 μm. **(D)** BDQ strongly enriched in PMN that are not only associated with the foamy macrophage-enriched areas. SEM of zoom 2 (left) and the distribution of secondary ions ^31^P^−^, ^79^Br^−^ and ^32^S^−^ (right). Overlaid image of SEM and secondary ions is depicted at the center. Demarcation of PMN is outlined by the white boundary line. Scale bar = 5 μm **(E)** Left panel: Quantitative analysis of BDQ associated with Mtb in BDQ treated and untreated mice. Center panel: Quantitative analysis of BDQ associated with LD in BDQ treated and untreated mice. Right panel: BDQ association to Mtb inside or outside LD in untreated vs BDQ-treated mice. Data show mean ± standard deviation. t-test adjusted for multiple comparisons. ns=non-significant; p value is as shown. At least 140 objects were counted from each treatment condition.

## Discussion

Altogether, we report the development of a correlative approach in tissue to define the subcellular localisation of antibiotics in infected cells within tissues. This multimodal imaging approach represents a powerful methodological advance to investigate if drugs reach their intracellular targets. Using this approach, we identified that in the lungs of *M. tuberculosis* infected mice, the antibiotic BDQ heterogeneously localised to intracellular bacteria and LD of foamy macrophages. We also found that BDQ significantly accumulated in specific cell types such as PMN, likely neutrophils, recruited into granulomatous lesions. Therefore, CLEIMiT enabled us to characterise the antibiotic distribution across multiple cell types, revealing multiple other niches of drug accumulation.

CLEIMiT is readily applicable to other drugs and not only for antibiotics or bromine-containing drugs but also any drugs detectable by nanoSIMS. Ion microscopy methods using nanoSIMS represents a good combination of spatial resolution and sensitivity to map drugs that contain elements other than Bromine that are low in the biological systems such as platinum [16], gold [17] and iodine [18]. What makes the approach more widely applicable is the nanoSIMS capability to detect stable isotope labelled molecules with high-resolution such as molecules that are labelled with ^2^H [19, 20], ^13^C [19, 21] and ^15^N [22, 23]. All of these stable isotopes can be used to label drugs of interest and mapped using the multiplexed imaging potential of CLEIMiT.

Importantly, in this study, we used a physiologically relevant treatment dose of BDQ. The detection limit of each elements or isotopes are different with nanoSIMS, and they are still not well documented in biological systems. However, multiple studies have demonstrated its capability to map drugs and other molecules with high resolution and sensitivity. From a pharmacokinetics/pharmacodynamics point of view, it would be important to define where and when the PMN internalise the antibiotic, since these cells are motile and actively recruited during lung inflammation. CLEIMiT also opens the possibility to define if other antibiotics currently used in the clinic are able to penetrate intracellular environments containing bacteria. Moreover, the combination of CLEIMiT with transgenic mice expressing specific fluorescent markers of cellular subtypes (e.g. myelocytic, endothelial, epithelial *etc*) will provide a suitable experimental setting to define in which cells antibiotics preferentially distribute and/or accumulate.

## Material and Methods

### Murine aerosol *M. tuberculosis* infection

*M. tuberculosis* H37Rv WT was kindly provided by Douglas Young (The Francis Crick Institute, UK). E2Crimson-Mtb was generated by transformation with pTEC19, a gift from Lalita Ramakrishnan (Addgene 30178). Bacteria were verified by sequencing and tested for the virulence-related lipids Phthiocerol dimycocerosates (PDIM) positivity. C3HeB/FeJ mice were bred under pathogen-free conditions at The Francis Crick Institute. Animal studies and breeding were approved by The Francis Crick Institute (London, UK) ethical committee and performed under UK Home Office project license PPL 70/8045. Infections were performed in the category 3 animal facility at The Francis Crick Institute. For aerosol infection, *M. tuberculosis* expressing E2crimson were grown to mid log-phase (OD_600_ of 0.6) in 7H9 (Sigma-Aldrich, M0178) supplemented with 10% albumin dextrose-catalase (BD Biosciences, 212351) and 0.05% TWEEN-80 (Sigma-Aldrich, P1754). An infection sample was prepared from this, to enable delivery of approximately 100 CFU/mouse lung using a modified Glas-Col aerosol infection system.

### Treatment with Bedaquiline

Three weeks after infection the treatment group was given 25mg/kg of Bedaquiline (BDQ, dissolved in 2-hydroxypropyl-β-cyclodextrin) (MedChemTronica, HY-14881) daily for 5 days via oral gavage while the control group were given only 2-hydroxypropyl-β-cyclodextrin. At the end of treatment, mice were euthanised by anaesthesia, then the lungs were either perfused with 10% neutral buffered formalin and excised **(Figure S1A)** or bacterial counts were determined by plating serial dilutions of homogenates on duplicate Middlebrook 7H11 (Sigma-Aldrich, M0428) containing OADC (BD Biosciences, 212240). Colonies were counted 2-3 weeks after incubation at 37°C. The data at each time point are the means of 5 mice/group +/– SEM **(Figure S1B)**. CFU/lung are calculated from the average of the duplicate multiplied by the volume of the dilution and the sample volume.

### CLEIMiT (Correlative Light, Electron and Ion Microscopy in Tissue)

#### Micro Computed tomography (μCT)

μCT imaging of whole lung: Whole lungs were incubated overnight at room temperature in 25% isotonic Lugol’s solution w/v for contrast then in 0.75 % low melting point agarose (LMA, 16520050, ThermoFisher Scientific) w/v in 200 mM HEPEs incubated at 37°C for 1h. Lungs were then set in 4% LMA and imaged using a Xradia 510 Versa 3D X-ray microscopes (Zeiss, Germany) with the following acquisition setting: 0.4X objective, pixel size = 7 μm, pixel binning of 2, source filter = LE1, Voltage = 40 kV, Wattage = 3.0 W. Tomogram reconstruction was carried out using the Zeiss Scout and Scan Software (Zeiss, Germany). Visualisation and fine measurements were taken from a 3D volume reconstruction using Zeiss XM3D viewer software (Zeiss, Germany).

μCT imaging of resin embedded tissue: A tissue slice was imaged using a Xradia 510 Versa 3D X-ray microscopes (Zeiss, Germany) with the following acquisition settings: 4X objective, pixel size = 2.8 μm, pixel binning of 2, source filter = LE2, Voltage = 40 kV, Wattage = 3.0 W. Tomogram reconstruction was carried out using the Zeiss Scout and Scan Software (Zeiss, Germany). Visualisation and fine measurements were taken from a 3D volume reconstruction using Zeiss’ XM3D viewer software (Zeiss, Germany). 3D measurements of the resin section were used to give precise co-ordinates for the location of the fluorescently imaged area in the resin block **(Figure S2B)** and to determine the precise angle of advance for the diamond knife when trimming the resin block.

#### Vibratome Sections

The lungs were separated into the four constituent lobes of the right lung (superior, middle, inferior, and post-caval) and left lung. Lobes were then embedded separately in 4% w/v LMA in 200 mM HEPES in agarose moulds (Sigma-Aldrich, E6032-1CS). The mould was placed on ice to cool and harden for sectioning. 100 μm sections were cut using a VT1200S fully automated vibrating blade microtome (Leica Biosystems, Germany). Upon calibration of the instrument, sections were cut at a cutting speed of 0.35mm/sec and an amplitude of 1.0 mm. Individual sections were sequentially removed and collected in a pre-labelled 12 well plate containing 1000 μl of 200 mM HEPEs buffer.

#### Fluorescence staining and imaging of mouse-lung sections

In a 24-well plate, 100μm lung slices were washed twice in 200 mM HEPES buffer then incubated for 20 min in a staining solution containing: 0.715 μM DAPI (4′,6-diamidino-2-phenylindole) (ThermoFisher Scientific D1306), 10 mg/L BODIPY 493/503 (4,4-difluoro-1,3,5,7,8-pentamethyl-4-bora-3a,4a-diaza-s-indacene) (Invitrogen D3922) in 200 mM HEPES buffer. Slices were washed twice with 200 mM HEPES and transferred to a glass slide and positioned to lie flat, unfolded across the surface. Excess buffer was removed, along with any remaining agarose and 10 μL DAKO fluorescent mounting medium (Agilent S3023) added. A cover glass (NA=1.5) was gently placed upon the tissue and the medium allowed to set. An inverted Leica TCS SP8 microscope running LAS X acquisition software with Navigator module (Leica Microsystems, Germany), equipped with 405 nm, Argon laser, 561 nm, 633 nm, and HyD detectors was used to image the tissue fluorescence with the following Lasers: 405nm (DAPI), 488nm (BODIPY) and 561nm (Mtb-E2Crimson). In the first instance, the entire tissue section was imaged with a tile scan using the 10x objective lens. Regions of interest (ROI) were then identified based upon areas of tissue showing high degrees of cellular infiltration, indicated by DAPI staining, and the accumulation of highly-lipid foamy cells, indicated by BODIPY staining. Selected ROI were then imaged at higher resolution using a 40x oil objective and z-stack. Voxel size was adjusted to half the theoretical limit of the lens in x and y and 0.5 μm in z. Fields of view were chosen to include cellular architecture such as airway passages as well as erythrocytes and vessels of the circulatory system, which appear as open space in the tissue, and can later be used as landmarks to help locate the ROI in downstream correlation. After imaging, slides were submerged in 200 mM HEPES and incubated at 4°C until the mounting medium dissolved and the tissue was released. Slices were then stored in 1.25 % glutaraldehyde (Sigma G5882), in 200 mM HEPES (Sigma-Aldrich H0887), pH 7.4 until embedding.

#### Resin embedding

Fluorescently imaged slices were processed for Scanning Electron Microscopy (SEM) and nanoscale secondary ion mass spectrometry (nanoSIMS) in a Biowave Pro (Pelco, USA) with use of microwave energy and vacuum. Samples (~0.3-0.4 mm^3^) were twice washed in HEPES (Sigma-Aldrich H0887) at 250 W for 40 s, post-fixed using a mixture of 2% osmium tetroxide (Taab O011) 1.5% potassium ferricyanide (Taab, P018) (v/v) at equal ratio for 14 min at 100 W power (with/without vacuum 20 “Hg at 2-min intervals). Samples were washed with distilled water twice on the bench and twice in the Biowave 250 W for 40 s, 1% thiocarbohydrazide (Sigma-Aldrich 223220) in distilled water (v/v) for 14 min at 100 W power (with/without vacuum 20 “Hg at 2 min intervals), washing cycle was repeated as before, then incubated with 2 % osmium tetroxide (Taab, O011) distilled water (w/v) for 14 min at 100 W power (with/without vacuum 20 “Hg at 2 min intervals). Samples were washed as before. Samples were stained with 1 % aqueous uranyl acetate (Agar scientific AGR1260A) in distilled water (w/v) for 14 min at 100 W power (with/without vacuum 20 “Hg at 2 min intervals) then washed using the same settings as before. Samples were dehydrated using a step-wise ethanol series of 50, 75, 90 and 100 %, then washed 4x in absolute acetone at 250 W for 40 s per step. Samples were infiltrated with a dilution series of 25, 50, 75, 100 % Durcupan ACM^®^ (Sigma-Aldrich 44610) (v/v) resin to acetone. Each step was for 3 min at 250 W power (with/without vacuum 20 “Hg at 30 s intervals). Samples were then cured for a minimum of 48 h at 60°C.

#### Resin block trimming

Referring to measurements from the 3D volume reconstruction, generated by μCT, the sample block was trimmed, coarsely by a razor blade then finely trimmed using a 35° ultrasonic, oscillating diamond knife (DiATOME, Switzerland) set at a cutting speed of 0.6 mm/s, a frequency set by automatic mode and a voltage of 6.0 V, on a ultramicrotome EM UC7 (Leica Microsystems, Germany) to remove all excess resin and tissue surrounding the ROI. Precise measurements, derived from the μCT reconstruction, were used to further cut into the tissue, to the depth corresponding with the fluorescent area previously imaged.

### Nanoscale secondary ion mass spectrometry (nanoSIMS)

The sections were imaged by SEM and nanoSIMS as previously described[10]. 500 nm sections were cut using ultramicrotome EM UC7 (Leica Microsystems, Germany) and mounted on 7 mm by 7 mm silicon wafers. Sections on silicon wafers were imaged using a FEI Verios SEM (Thermo Fisher Scientific, USA) with a 1 kV beam with the current at 200 pA. The same sections were then coated with 5 nm gold and transferred to a nanoSIMS 50L instrument (CAMECA, France). The regions that were imaged by SEM were identified using the optical microscope in the nanoSIMS. A focused ^133^Cs^+^ beam was used as the primary ion beam to bombard the sample; secondary ions (^12^C^−^, ^12^C^14^N^−^, ^79^Br^−^, ^32^S^−^ and ^31^P^−^) and secondary electrons were collected. A high primary beam current of ∼1.2 nA was used to scan the sections to remove the gold coating and implant ^133^Cs^+^ to reach a dose of 1×10^17^ ions/cm^2^ at the steady state of secondary ions collected. Identified regions of interest were imaged with a ~3.5 pA beam current and a total dwell time of 10 ms/pixel. Scans of 512 × 512 pixels were obtained.

#### Image alignment

Tissue derived micrograph and nanoSIMS/micrograph correlation: ion and fluorescent images were aligned to EM micrographs with Icy 2.0.3.0 software (Institut Pasteur, France), using the ec-CLEM Version 1.0.1.5 plugin. No less than 10 independent fiducials were chosen per alignment for 2D image registration. When the fiducial registration error was greater than the predicted registration error, a non-rigid transformation (a nonlinear transformation based on spline interpolation, after an initial rigid transformation) was applied as previously described[24].

#### Quantification and Statistical analysis

Ion quantification: secondary Ion signal intensities were quantified in ImageJ with the OpenMIMS v3.0.5 plugin.

Quantification of BDQ within bacteria: bacteria (_total_n = 472) were manually outlined with the assistance of SEM images and the ^31^P^−^ signal. Ratio values (^79^Br^−^/^12^C^14^N^−^) for bacteria were divided by the area of their respective ROI to give mean normalised pixel intensity in arbitrary units (AU) for each condition. Mean normalised pixel intensity in arbitrary units (AU) per ROI was plotted against condition using Graphpad Prism 8 software and two-tailed p-value was determined by an unpaired, non-parametric Mann-Whitney U test to assess statistical significance.

Quantification of BDQ in lipid droplets: lipid droplets (_total_n = 1404) were outlined using the (^32^S^−^/1) ratio value. The resulting ratio image was summed and processed with a gaussian blur filter (sigma radius = 2 pixels). A threshold was applied to mask the image. ROIs were identified by particle analysis, and verified by comparison with the respective SEM image. Masked areas were overlaid to the ^79^Br^−^/^12^C^14^N^−^ ratio image of the same area of tissue. ROIs with less than 5 pixels in size were excluded from the analysis. Mean normalised pixel intensity in arbitrary units (AU) per ROI was plotted against condition using GraphPad Prism 8 software. Two-tailed p-value was determined by an unpaired, non-parametric Mann-Whitney U test to assess statistical significance.

Quantification of BDQ in bacteria inside LD: bacteria (_total_n = 282) were manually outlined and localisation defined to be either inside lipid droplets (inLD) or outside lipid droplets (outLD) with the assistance of SEM images and the ^31^P^−^ and ^32^S^−^ signal. Ratio values (^79^Br^−^/^12^C^14^N^−^) for bacteria were divided by the area of their respective ROI to give mean normalised pixel intensity in arbitrary units (AU) for each condition. Two-tailed p-values were determined by an unpaired, non-parametric Kruskal-Wallis test with Dunn’s correction to assess statistical significance.

Quantification of BDQ in PMN: Polymorphonuclear cells (_total_n = 22) were manually outlined with the assistance of SEM images and the ^12^C^14^N^−^ signal. Ratio values (^79^Br^−^/^12^C^14^N^−^) for PMN were divided by the area of their respective ROI to give mean normalised pixel intensity in arbitrary units (AU) for each condition. Mean normalised pixel intensity in arbitrary units (AU) per ROI was plotted against condition using Graphpad Prism 8 software. Two-tailed p-value was determined by an unpaired, non-parametric Mann-Whitney U test to access statistical significance.

## Acknowledgements

We thank Elliott Bernard (The Francis Crick Institute) for making the Mtb fluorescent strain used in this work and Gareth Griffiths (University of Oslo) for useful suggestions on the manuscript. We are also grateful to the Advanced Light Microscopy STP, the Electron Microscopy STP and Biological Research Facility at the Crick for their support in various aspects of the work. We thank Paul Guagliardo, Jeremy Bougoure and Alexandra Suvorova (Centre for Microscopy, Characterisation and Analysis, University of Western Australia) for their support for nanoSIMS and SEM imaging. This work was supported by the Francis Crick Institute (to MGG), which receives its core funding from Cancer Research UK (FC001092), the UK Medical Research Council (FC001092), and the Wellcome Trust (FC001092). HJ is supported by an Australian Research Council Discovery Early Career Researcher Award.

## Author Contributions

MGG and HJ conceived and supervised the project. AF, DJG, AR and HJ performed the experiments. AF performed the sectioning, EM sample preparation and the full correlative approach. DJG performed the fluorescence analysis. HJ performed the SEM and nanoSIMS. AR carried out the mouse infections, CFU analysis and tissue collection. AF, DJG and HJ analysed the data. MGG wrote the manuscript and prepared the figures with input from AF. All authors discussed the results and implications and commented on the manuscript at all stages.

## Supplementary information

**Supplementary Figure 1. Overview of the infection model and treatment.**

(A) Diagram of the infection and treatment experimental setting.

(B) Colony forming units (CFU) in the lungs of mice at day zero of infection (inoculum) and treated with either BDQ or vehicle.

(C) Micro Computed tomography (μCT) of whole lung showing granulomatous lesions.

**Supplementary Figure 2. Strategy for correlative light (fluorescence), electron (SEM) and ion (nanoSIMS) microscopy.**

(A) Diagram of the sectioning strategy for correlation, including the different imaging modalities and thickness. A^o^ represents the need to calculate and adjust the angle of incidence of the diamond knife with the resin block so as to achieve parallel sectioning from the surface of the tissue, through the entire block.

(B) Correlation between fluorescent ROI and μCT of resin embedded section and positioning localisation of the SEM section for correlation between nanoSIMS/SEM and fluorescence. Middle panels show different orthogonal slices of the 3D section used to localise area, and calculate angle and depth for further sectioning. The intersecting lines indicate the precise location of the target cell within the resin embedded section. Distances which are used to calculate angles are shown in blue numbers. These localisations were used to zoom in the ROI (zoom) and obtain the SEM image corresponding to the fluorescent image in the upper panel.

**Supplementary Figure 3. BDQ accumulates heterogeneously in LD and Mtb, with particularly high levels in infected foamy macrophages.**

**(A)** Fluorescent microscopy of a Region of Interest (ROI) as in Figure 1: (i) cellular infiltration and the accumulation of BODIPY (green) positive cells. Cells infected with Mtb-E2Crimson (red) are also visible throughout this region. Scale bar = 100 μm. (ii) Zoomed in image showing the selected foamy cell infected with Mtb-E2Crimson (red) from (i) for correlative analysis. Scale bar = 15 μm. Lower panels show the ion microscopy for the selected cell including ^12^C^14^N^−^, ^31^P^−^, ^79^Br^−^ and ^32^S^−^. Compass indicates the orientations of secondary ion images with regard to the fluorescent image above.

**(B)** Correlative light, electron and ion microscopy in tissue (CLEIMiT): Right, a correlated image overlaying the ^79^Br^−^ and ^31^P^−^ signals with the SEM image. Center, the corresponding SEM image of the infected foamy cell. Left, a correlated image overlaying fluorescent signal from BODIPY (green) and Mtb-E2Crimson (red) against the SEM image. Scale bar = 5 μm.

**Supplementary Figure 4. Quantitative analysis workflow of BDQ levels in Mtb and LD**

(A) Masking of Mtb profiles aided by the combination of the SEM profiles and ^31^P^−^ signal and measurement of the ^79^Br^−^ signal associated with Mtb (see methods). Scale bar = 5 μm.

(B) Masking of LD profiles aided by the combination of the SEM profiles and ^32^S^−^ signal and measurement of the ^79^Br^−^ signal associated with LD (see methods). Scale bar = 5 μm.

**Supplementary Figure 5. Bromine signal from PMN is primarily associated with BDQ**

(A) Representative SEM/nanoSIMS correlated images of PMN in granulomatous lesions in lungs of mice treated with vehicle (untreated) and BDQ (treated).

(B) Quantitative analysis of ^79^Br^−^ signal in PMN. Data show mean ± standard deviation. t-test adjusted for multiple comparisons. ns=non-significant; p value is as shown. A total of 22 PMN were counted. Scale bar = 5 μm.

**Movie S1:**μCT of infected lungs showing lesions.

**Movie S2**: Ion images and intracellular localisation of BDQ in different cell types within granulomatous lesions.

